# COMPARATIVE PHYTOCHEMICAL ANALYSIS OF TWO ALPINIA SPECIES FROM ZINGEBERACEAE FAMILY

**DOI:** 10.1101/2023.10.25.564021

**Authors:** N. Rabita, K. Palanisamy

## Abstract

The traditional applications of the rhizomes of two Alpinia species, *Alpinia galanga* and *Alpinia nigra*, both of which belong to the Zingiberaceae family, are widely recognized. Alpinia has been concentrating on its biological defenses against terrible diseases including cancer and devastating viral infections. In order to develop the diagnostic keys for these significant medications based on the phytochemical analysis, the current research work was done to conduct a detailed comparative phytochemical investigation of the two species. *Alpinia galanga* and *Alpinia nigra* were the focus of this study’s evaluation of their phytochemical components. Alkaloids, saponins, quinones, terpenoids, steroids, tannins, flavonoids, cardiac glycosides, and coumarins were all investigated in our experiments to see if they were present within the two species. The primary chemical compounds found in *Alpinia galanga* and *Alpinia nigra* that may be significant from a pharmacological perspective are highlighted in this article. A qualitative phytochemical study of plant extracts revealed that most of the substances, such as quinones, terpenoids, tannins, steroids, flavonoids, cardiac glycosides, and coumarins, were present.

## Introduction

Medicinal plants are the most abundant natural sources of pharmaceuticals used in conventional and traditional treatment. Since a few decades ago, there has been an increase in interest in discovering the therapeutic properties of various plants. This is likely due to their potential to be sources of powerful pharmacological activities, user-friendliness, economic feasibility, and low toxicity. a wide range of medicinal plants that contain molecules that can scavenge free radicals, including vitamins, polyphenolic compounds, terpenoids, tannins, coumarins, alkaloids, amines, and other metabolites with strong antioxidant activity. It is therefore imperative to make ongoing efforts to find new traditional medicine sources and to check the existing ones for new therapeutic indications.

There are 230 species in the Zingiberaceae genus *Alpinia* Roxb., of which roughly 50 are found in east and southwest China. It is the largest, most geographically diverse, and most taxonomically complicated genus. The majority of them are found in Asia’s tropical and subtropical regions, including India, Malaysia, China, and Japan. In several nations, including China, Japan, and India, plants in this genus have been used as traditional foods, medicines, and spices. *A. galanga* (L.) Willd is an essential component of curries and has been widely used as a flavoring in the preparation of meats and soups in Southeast Asia as well as in the preparation of beverages in Europe. Other examples include *A. vittata, A. purpurata* (Vieill.) K. Schum, *A. calcarata* Rosc, and *A. zerumbet*, which are grown for their ornamental qualities. The biological defenses of the Alpinia genus against deadly diseases like cancer and fatal viral infections have recently been the focus of research.

Greater galangal, also known as *Alpinia galanga*, is a significant perennial herb from the Zingiberaceae family that is grown in countries including India, China, Thailand, Malaysia, and Indonesia. It is also known as Kulanjan in Hindi and Kanghoo in Manipuri. Alpinia galanga rhizomes are used medicinally for the treatment of conditions such as neuroprotection, antileishmanial, antiallergic, immune-stimulating, hypoglycemic, antimicrobial antitumor, antifungal, antioxidant and inhibitors of nitric oxide synthesis. *Alpinia galanga* is primarily used by traditional South Indian ayurvedic and Siddha medical practitioners to treat a variety of illnesses, including diabetes mellitus. There have been claims that the rhizome extract of *Alpinia galanga* contains a number of medicinally active compounds with antitumor, antioxidant, antifungal, antibacterial, gastroprotective, hypoglycaemic, hypolipidemic, and anti-inflammatory properties.

The Zingiberaceae family member *Alpinia nigra* B.L. Burtt, also known as Jongly Ada or Tara in Bengali, and generally known as “Noh Kala” in Thailand, is also widely cultivated in China’s Yunnan and Hainan provinces as well as other Southeast Asian nations. Other names for this fragrant, rhizomatous herb are Kala, False galangal, Greater galangal, Black-Fruited, and False galangal. Natives of Tripura, India, use the aqueous juice of *Alpinia nigra* shoots to treat intestinal parasite illness. In addition to being a vegetable that may be eaten, it is also used in traditional medicines to cure insect bites, gastrointestinal illness, and dyspepsia. Native communities in Tripura, Northeast India, have long used the shoot of this plant, consuming the raw juice of the green shoot due to its rumored anthelmintic effects. Thailand also consumes the rhizomes of this plant as veggies. Galangal, curcuma, and zinger have close kinship with the rhizomes of *Alpinia nigra*. Two flavone glycosides, astragalin and kaempferol-3-O-glucuronide, were discovered during a prior phytochemical inquiry. These compounds have been shown to have a variety of biological properties, including antibacterial, antioxidant, antiprotozoal, shoots have all been shown to have antibacterial, anti-inflammatory, and anthelmintic properties in recent investigations.

## Material and methods

### Collection of plant materials

The fresh rhizomes of plant material used in the present study namely *Alpinia galanga* and *Alpinia nigra* was collected in the month of March and April from Bishnupur District, Manipur, North East, India.

### Preparation of plant material

The fresh rhizomes were first cleaned with running water to get rid of any dirt, and then distilled water was used for the last cleaning. The cleaned rhizomes were next allowed to air dry, thinly slice, and dry at room temperature before being ground into a coarse powder. The resulting fine powder was then kept at 4ºC in an airtight container for future experimental usage.

### Preparation of rhizome extract

The collected rhizome was shade dried, pulverized, and extracted for eight hours using a Soxhlet apparatus with hexane, chloroform, acetone, methanol, and water. The crude extract’s solvent was removed from the extract using a rotary vacuum evaporator at a temperature of 60ºC until the solvent had fully evaporated. In order to conduct further research, the collected residue was kept in the refrigerator.

### Yield of extract (%)

The weight of crude extract produced after extraction was divided by the weight of plant powder weighed before to extraction, and the result was multiplied by 100 to determine the percentage of extraction yield (%).

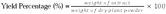

### Preliminary Phytochemical Screening

In order to identify the phytochemical constituents, a preliminary phytochemical screening was done for the presence of secondary metabolites, particularly alkaloids, saponin, quinones, terpenoids, carbohydrate, steroids, tannin, amino acids and protein, flavonoids, cardiac glycosides, and coumarin.

### Test for alkaloids

2 drops of Mayer’s reagent were added to a few ml of filtrate. Alkaloids are present when a precipitate with a creamy white color form.

### Test for saponin

The extract 0.5g was mixed with five ml of distilled water. Saponins are present when there is formation of foaming (creation of creamy-looking, tiny bubbles that last for about 15 minutes).

### Test for quinone

1ml of the extract was mixed with 1ml of concentrated H_2_SO_4_; the formation of a red color indicates the presence of quinones.

### Test for Terpenoids

When three drops of chloroform and two drops of concentrated H_2_SO_4_ were added to the extract, the lower portion took on a yellow or reddish-brown color, which indicates the presence of terpenoids.

### Test for carbohydrate

Few ml of plant extract were added to 5 ml of Benedict’s reagent, heated for 2 minutes, and then cooled. The presence of carbohydrates is shown by the production of a red precipitate.

### Test for Steroids

Chloroform and concentrated H_2_SO_4_ were added to 2 ml of rhizome extract. Reddish-brown ring formation indicates the presence of steroids.

### Test for Tannin

When a few ml of potassium dichromate is added to the plant extract, yellow precipitate forms, indicates the presence of tannins.

### Test for Amino acid and protein

In a test tube, 4% NaOH and 1% CuSO_4_ were combined with the extract solution. When proteins were present, the color of the solution changed to violet or pink.

### Test for Flavonoids

To 2 ml of the extract, 2 drops of sodium hydroxide (NaOH) were added. When a few drops of diluted HCl were added, the intense yellow color that initially appeared eventually faded, showing that flavonoids were present.

### Test for Cardiac Glycosides

A few ml of glacial acetic acid, ferric chloride, and concentrated H_2_SO_4_ were added to the rhizome extract. The presence of cardiac glycosides is indicated by the color of bluish green.

### Test for Coumarins

1 mL of rhizome extract was mixed with 1 mL of 10% NaOH. The formation of yellow color denotes the presence of coumarins.

## RESULTS AND DISCUSSION

### Extractive yield

The percentage yield of *Alpinia galanga* and *Alpinia nigra* in different extracts was shown in Table 1. The findings show that, in contrast to other extracts, methanol extract has a higher yield of extract (11.07% and 8.5%, respectively), whereas chloroform extract has a lower yield, revealing that chloroform has a poor ability to extract *Alpinia galanga* and *Alpinia nigra*.

**Table 1:**
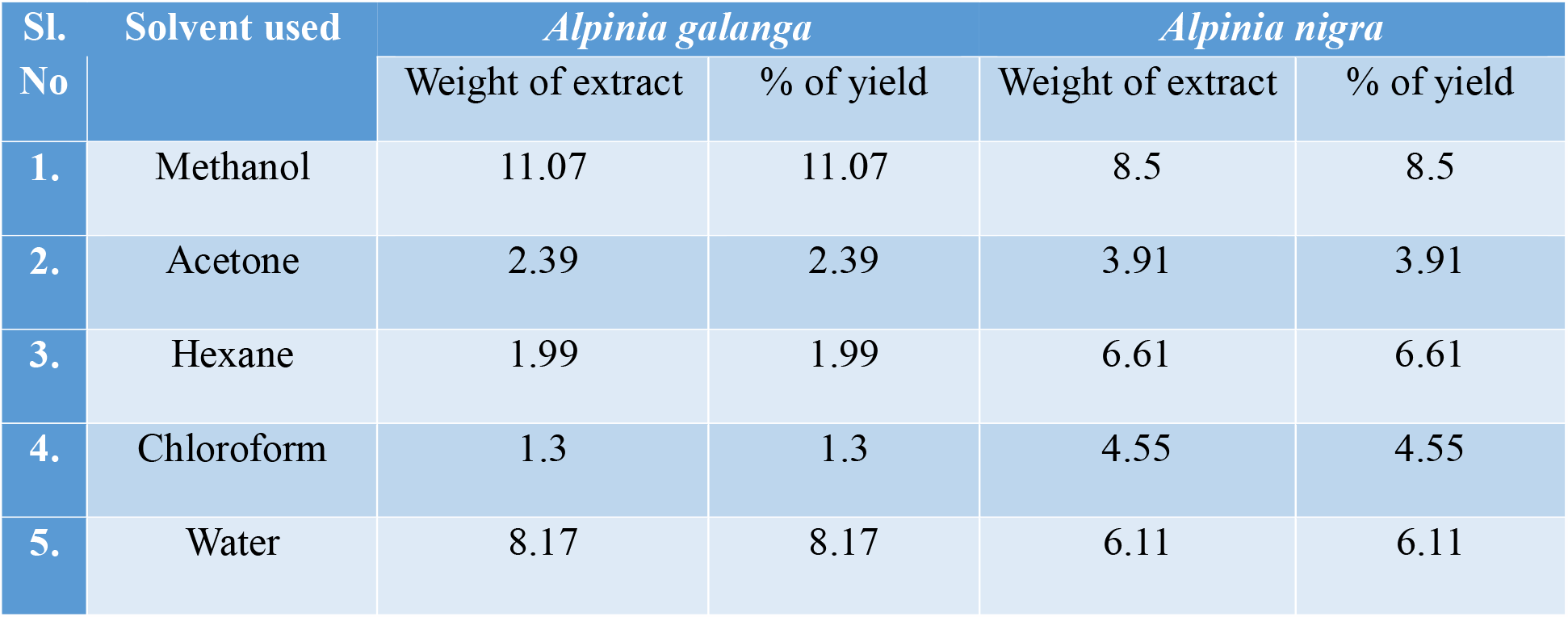
Percentage yield of different extracts of *Alpinia galanga* and *Alpinia nigra*.

### Phytochemical analysis

A preliminary phytochemical analysis of eleven bioactive components from various extracts of *Alpinia galanga* and *Alpinia nigra* was performed, and the findings are shown in Tables 2 and 3, respectively. *Alpinia galanga* has quinones, terpenoids, steroids, tannins, flavonoids, cardiac glycosides, and coumarins, whereas *Alpinia nigra* includes alkaloids, saponin, quinones, terpenoids, steroids, tannins, flavonoids, and cardiac glycosides. Protein and amino acids did not exhibit any evidence that they were present in any of the various solvent extracts, as shown in the table. The presence of the greatest number of phytochemical components was found in the methanol extract, which was one of the five different *Alpinia galanga* and *Alpinia nigra* extracts. Depending on the solvent medium employed for extraction, phytochemicals may be present or absent. All of the phytochemicals examined were present in the different solvent extracts of *Alpinia galanga* and *Alpinia nigra*. The most prevalent phytochemicals in both plants were discovered to be quinones, terpenoids, tannins, steroids, flavonoids, cardiac glycosides, and coumarins. both results imply that both plants could be a viable source of bioactive substances with medicinal potentials, and both plants displayed comparable phytochemicals.

**Table 2:** Phytochemical constituents present in different extracts of *Alpinia galanga*.

| Sl. No. | Name of phytochemical compounds | Rhizome extracts of *Alpinia galanga* |  |  |  |  |
| --- | --- | --- | --- | --- | --- | --- |
|  |  | Methanol | Acetone | Hexane | Chloroform | Water |
| 1. | Alkaloids | + | - | + | + | - |
| 2. | Saponins | - | - | - | + | + |
| 3. | Quinones | + | + | + | + | - |
| 4. | Terpenoids | + | + | + | + | - |
| 5. | Carbohydrates | + | - | - | - | - |
| 6. | Steroids | + | + | - | + | - |
| 7. | Tannins | + | + | + | + | + |
| 8. | Amino acid and protein | - | - | - | - | - |
| 9. | Flavonoids | + | + | + | + | - |
| 10. | Cardiac Glycosides | + | + | + | - | - |
| 11. | Coumarins | + | + | + | + | - |

(+) indicates present, (-) indicates absent.

**Table 3:**
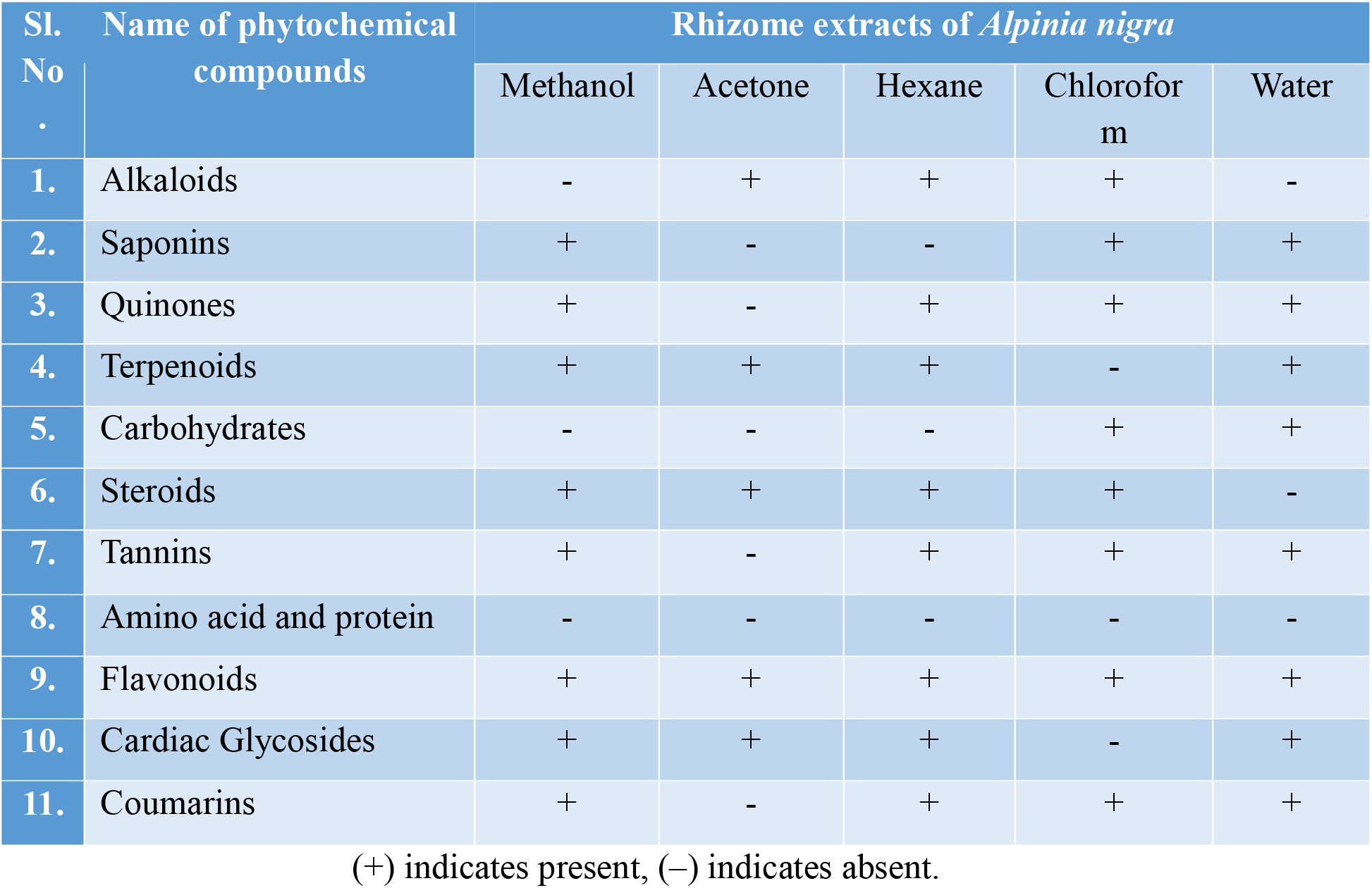
Phytochemical constituents present in different extracts of *Alpinia nigra*.

| Sl. No. | Name of phytochemical compounds | Rhizome extracts of *Alpinia nigra* |  |  |  |  |
| --- | --- | --- | --- | --- | --- | --- |
|  |  | Methanol | Acetone | Hexane | Chloroform | Water |
| 1. | Alkaloids | - | + | + | + | - |
| 2. | Saponins | + | - | - | + | + |
| 3. | Quinones | + | - | + | + | + |
| 4. | Terpenoids | + | + | + | - | + |
| 5. | Carbohydrates | - | - | - | + | + |
| 6. | Steroids | + | + | + | + | - |
| 7. | Tannins | + | - | + | + | + |
| 8. | Amino acid and protein | - | - | - | - | - |
| 9. | Flavonoids | + | + | + | + | + |
| 10. | Cardiac Glycosides | + | + | + | - | + |
| 11. | Coumarins | + | - | + | + | + |

(+) indicates present, (-) indicates absent.

The understanding of these plants phytochemical components will be useful for discovering low-cost, highly effective herbal folklore medicines in the future. Additional research is needed to isolate the active components in these extracts in order to clarify their medicinal potential and create a potentially useful pharmacological molecule for treating oxidative stress and related illnesses. In order to treat diabetics and other conditions linked to oxidative stress, this will aid in the development of new herbal medicines that are efficient, secure, and reasonably priced. The plant can be investigated further against different ailments to determine its potential efficacy and may be a source of chemically intriguing and physiologically significant medication possibilities.

**Fig 1:**
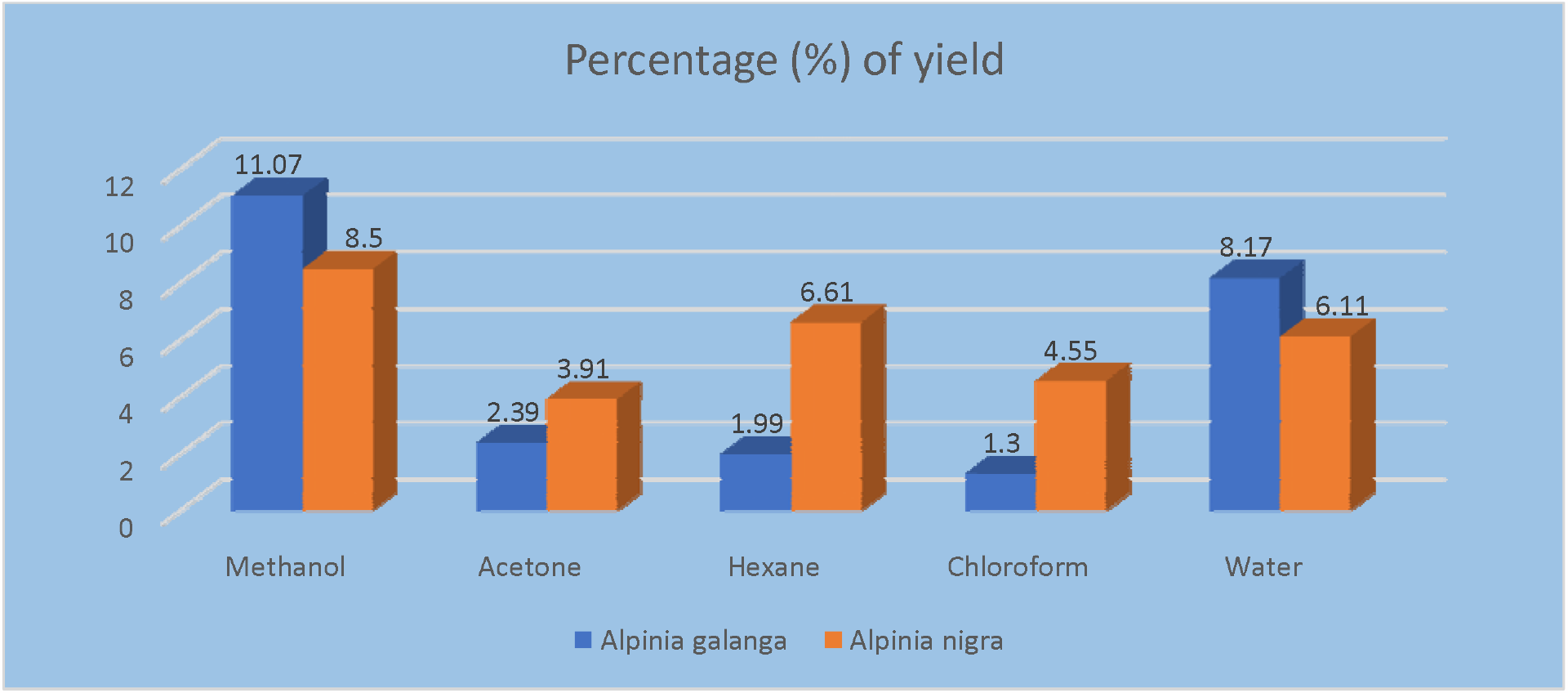
Percentage (%) of yield extract of *A. galanga* and *A. nigra* in different solvent.

**Fig 2:**
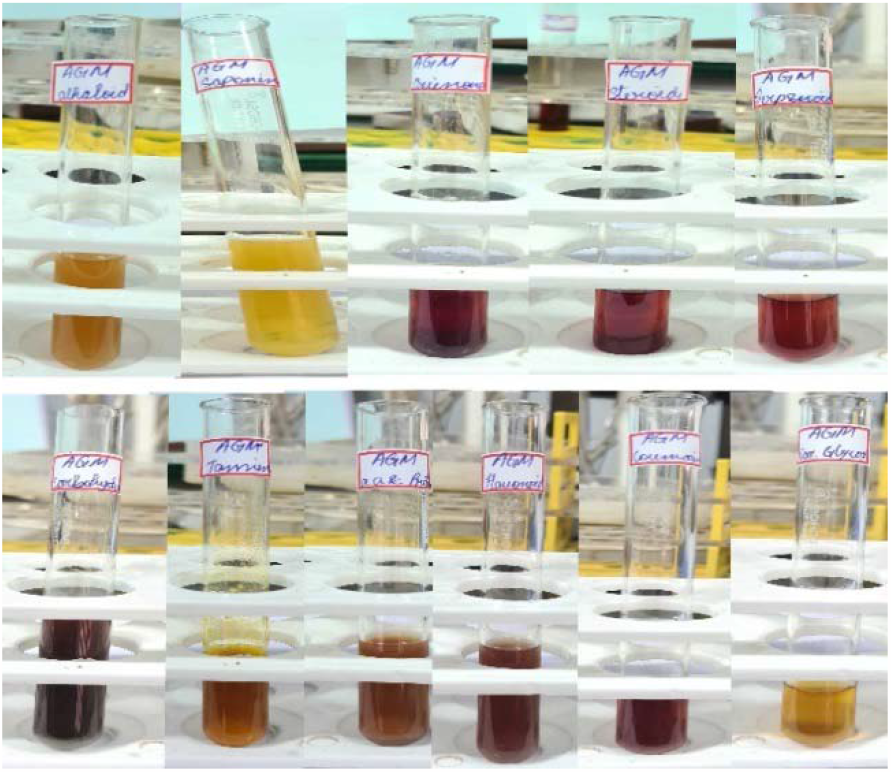
*Alpinia galanga* methanol extract.

**Fig 3:**
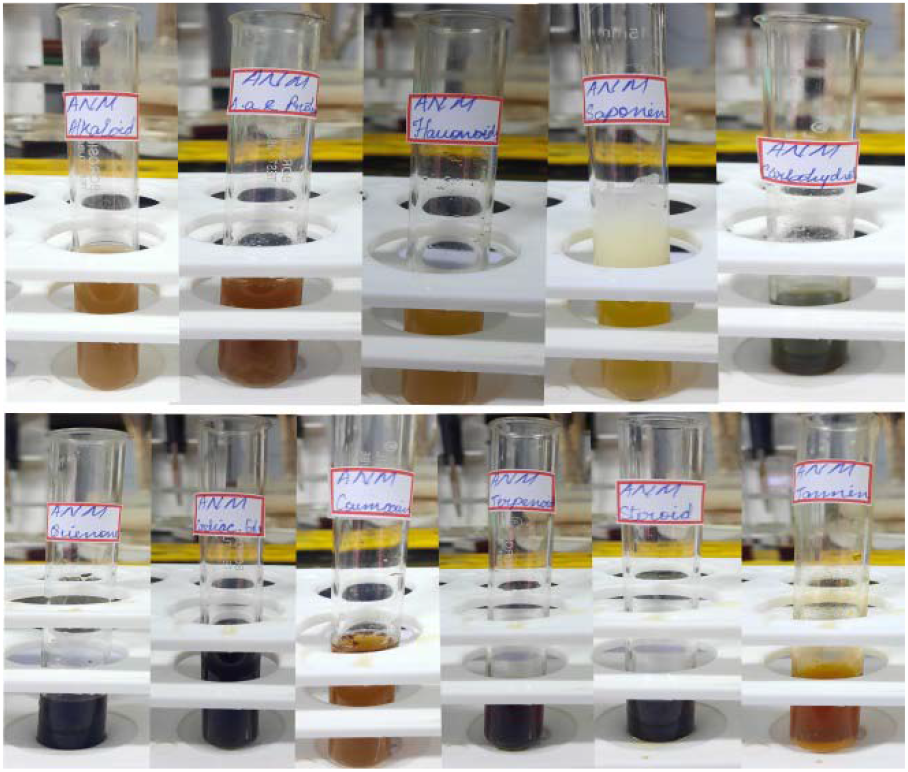
*Alpinia nigra* methanol extract.

**Fig 4:**
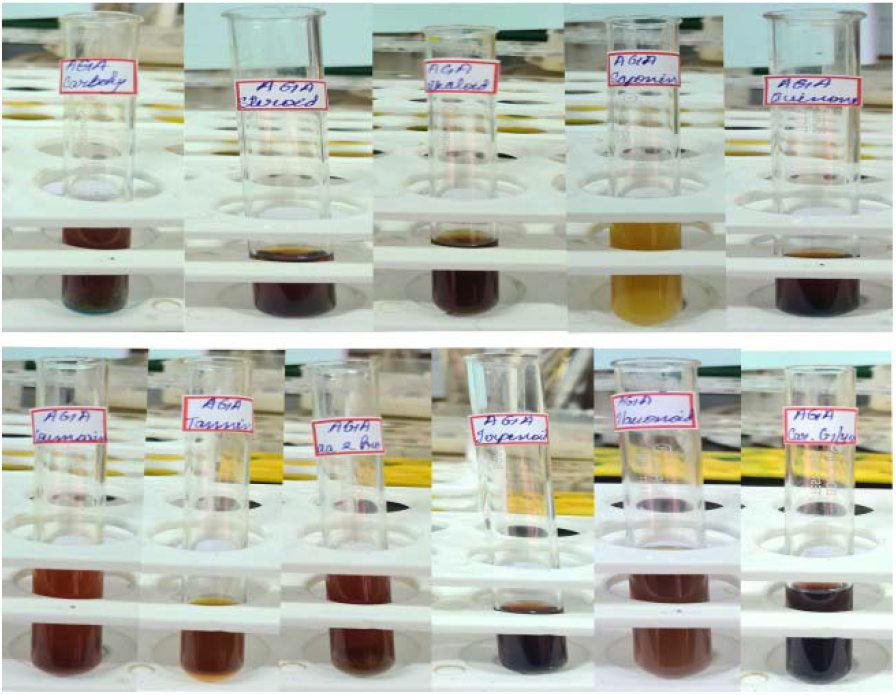
*Alpinia galanga* acetone extract.

**Fig 5:**
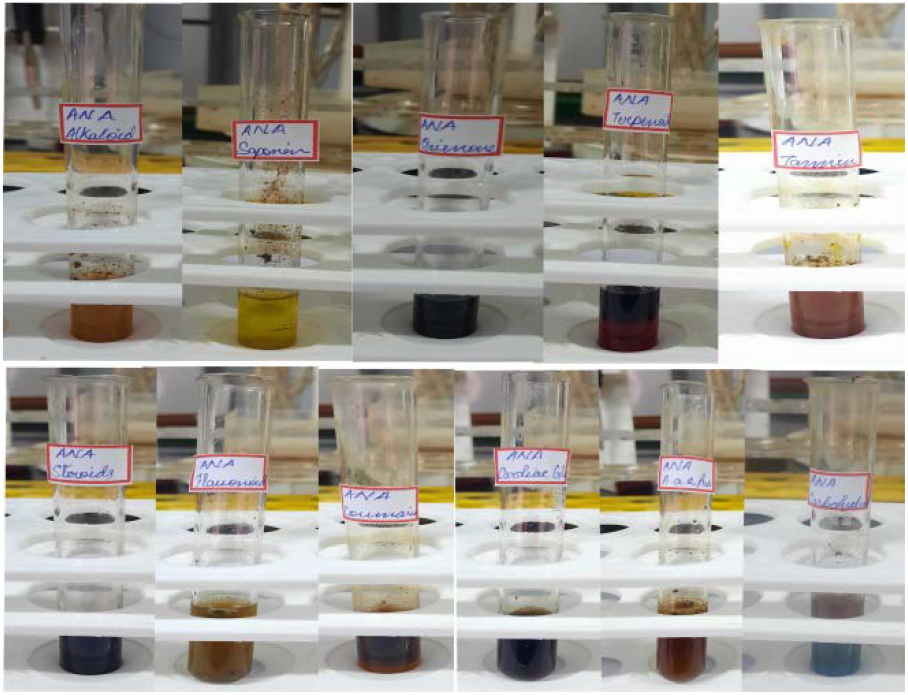
*Alpinia nigra* acetone extract.

**Fig 6:**
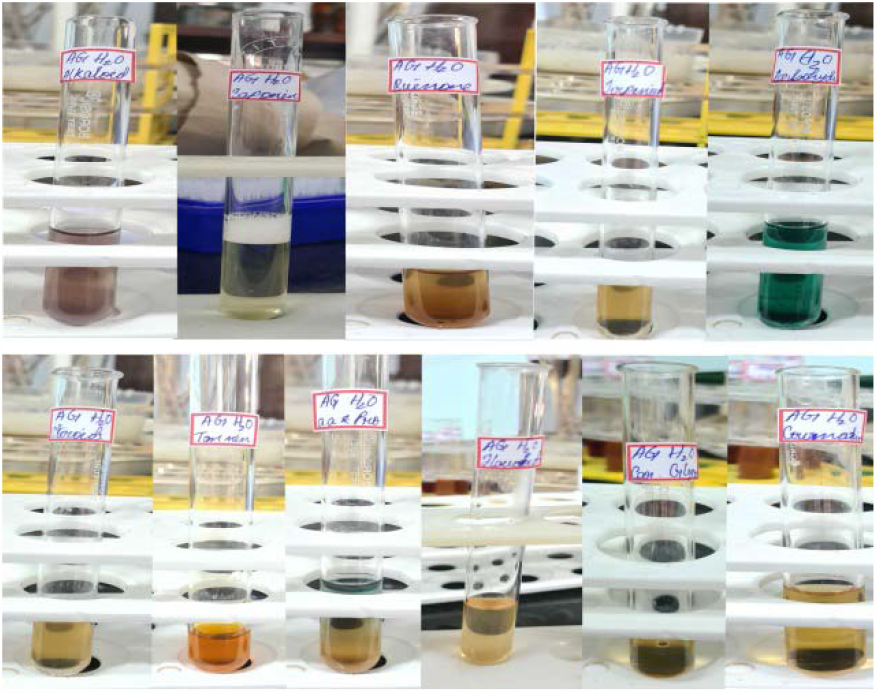
*Alpinia galanga* water extract.

**Fig 7:**
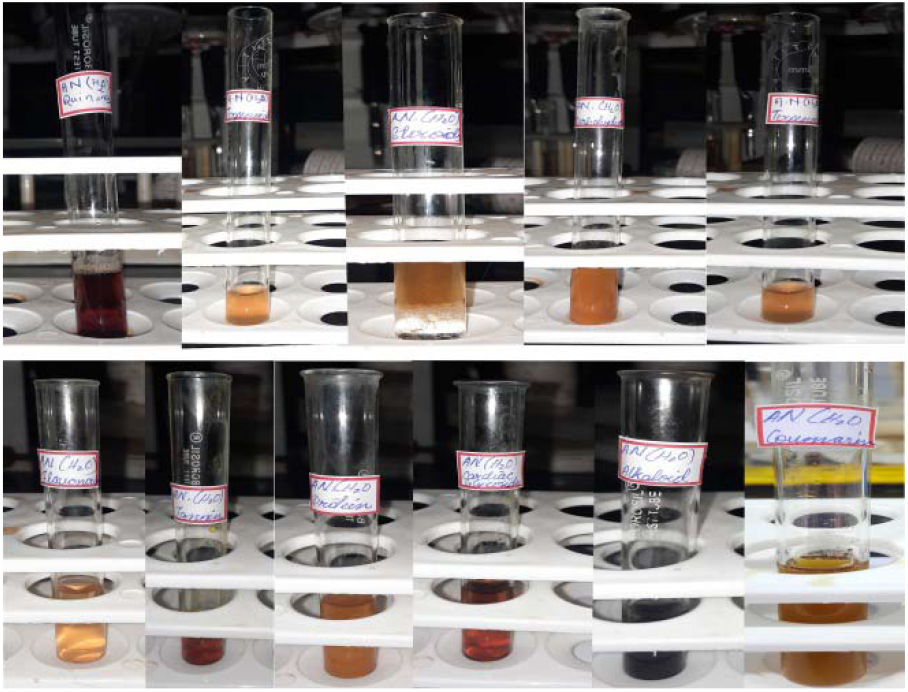
*Alpinia nigra* water extract.

**Fig 8:**
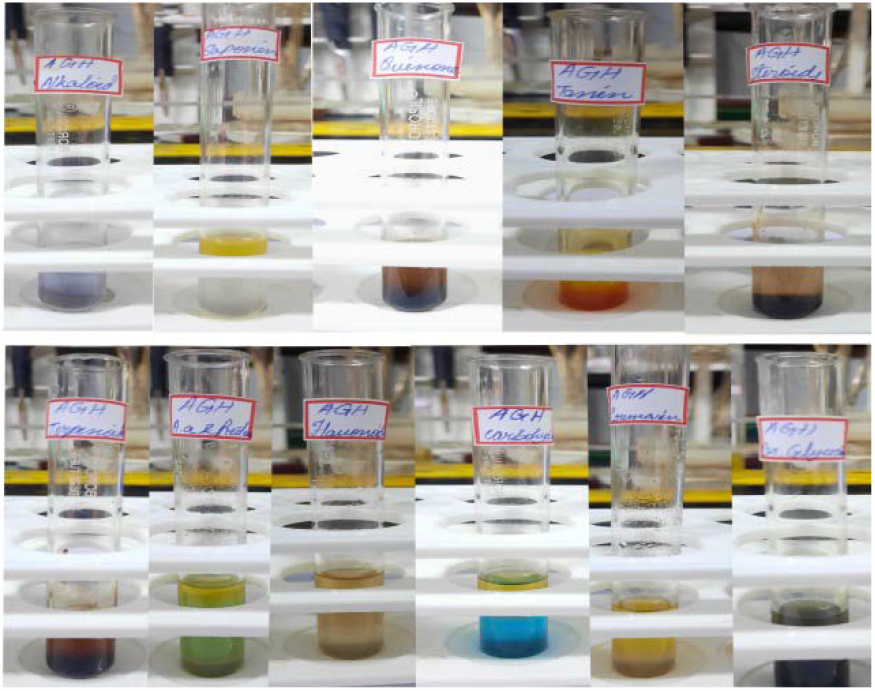
*Alpinia galanga* hexane extract.

**Fig 9:**
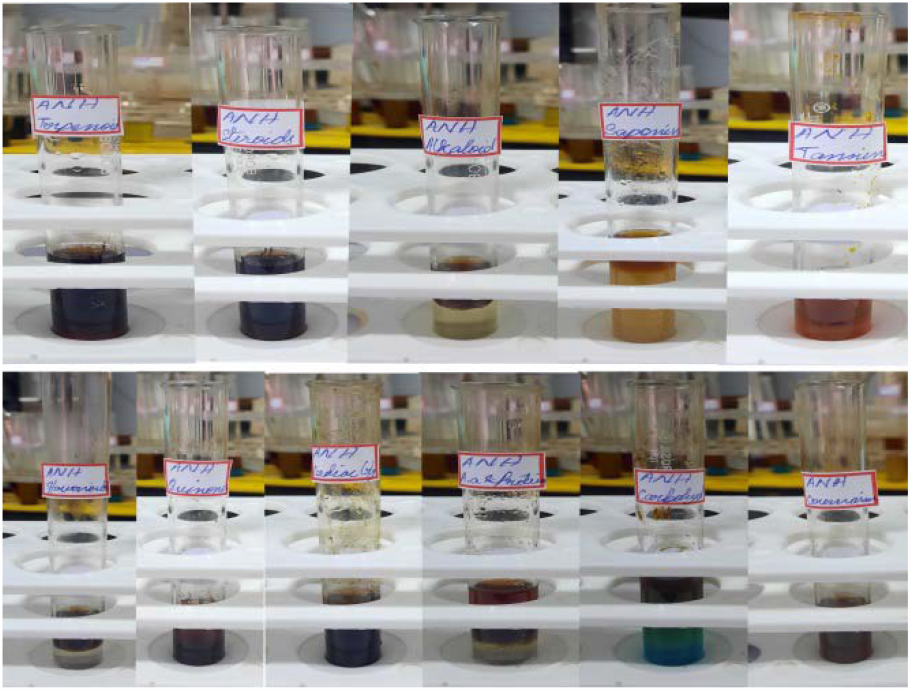
Alpinia nigra hexane extract.

**Fig 10.**
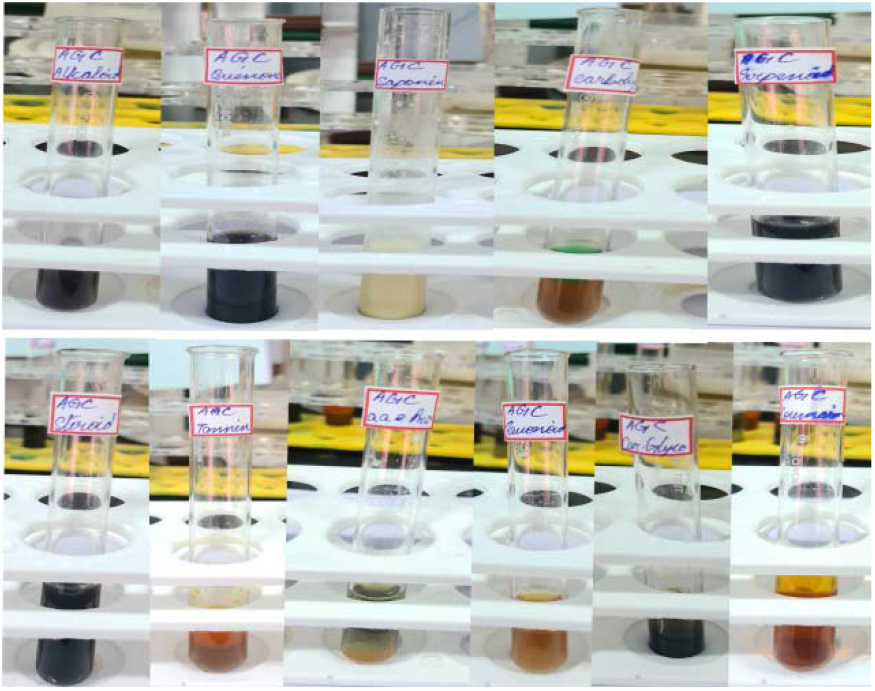
*Alpinia galanga* chloroform extract.

**Fig 11.**
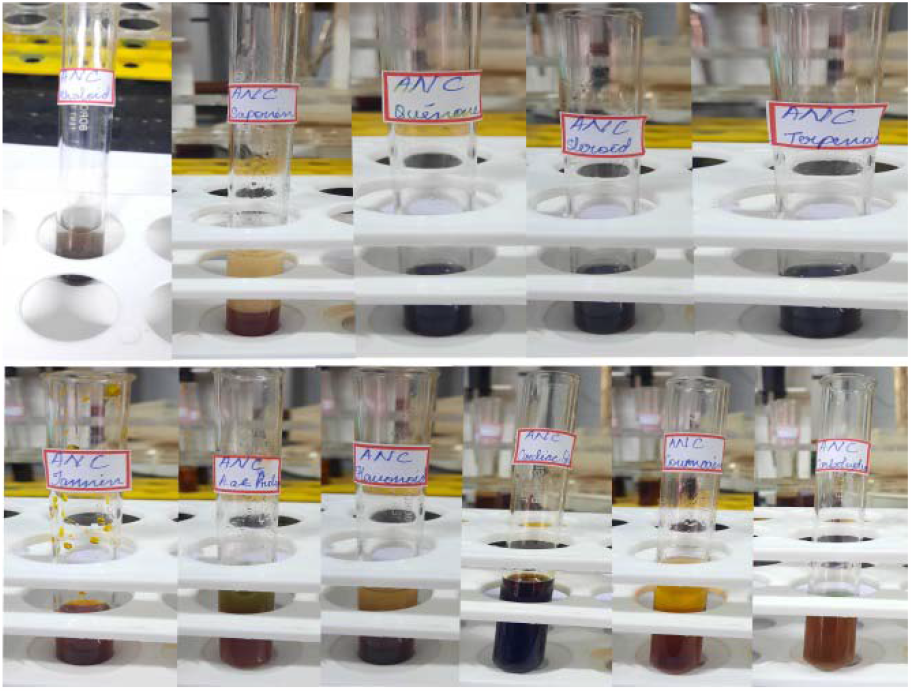
*Alpinia nigra* chloroform extract.

## Conclusion

From the results and discussion, study concluded that Quinones, terpenoids, tannins, steroids, flavonoids, cardiac glycosides, and coumarins were found to be the most abundant phytochemicals in the two Alpinia species. Both findings suggest that the two Alpinia species may represent potential sources of bioactive compounds with therapeutic potentials, both plants displayed comparable phytochemicals.

## CRediT authorship contribution statement

**N. Rabita:** Conceptualization, Investigation, Validation, Data curation, Visualization, Writing – review & editing. **K. Palanisamy:** Guiding.

## Acknowledgement

The authors were thankful to the Department of Botany, Annamalai University, Annamalai Nagar for providing necessary laboratory facilities to carry out present research work.

